# The role of untuned neurons in sensory information coding

**DOI:** 10.1101/134379

**Authors:** Joel Zylberberg

## Abstract

To study sensory representations, neuroscientists record neural activities while presenting different stimuli to the animal. From these data, we identify neurons whose activities depend systematically on each aspect of the stimulus. These neurons are said to be “tuned” to that stimulus feature. It is typically assumed that these tuned neurons represent the stimulus feature in their firing, whereas any “untuned” neurons do not contribute to its representation. Recent experimental work questioned this assumption, showing that in some circumstances, neurons that are untuned to a particular stimulus feature *can* contribute to its representation. These findings suggest that, by ignoring untuned neurons, our understanding of population coding might be incomplete. At the same time, several key questions remain unanswered: Are the impacts of untuned neurons on population coding due to weak tuning that is nevertheless below the threshold the experimenters set for calling neurons tuned (vs untuned)? Do these effects hold for different population sizes and/or correlation structures? And could neural circuit function ever benefit from having some untuned neurons vs having all neurons be tuned to the stimulus? Using theoretical calculations and analyses of *in vivo* neural data, I answer those questions by: a) showing how, in the presence of correlated variability, untuned neurons can enhance sensory information coding, for a variety of population sizes and correlation structures; b) demonstrating that this effect does not rely on weak tuning; and c) identifying conditions under which the neural code can be made more informative by replacing some of the tuned neurons with untuned ones. These conditions specify when there is a functional benefit to having untuned neurons.

**Author Summary:** In the visual system, most neurons’ firing rates are tuned to various aspects of the stimulus (motion, contrast, etc.). For each stimulus feature, however some neurons appear to be untuned: their firing rates do not depend on that stimulus feature. Previous work on information coding in neural populations ignored untuned neurons, assuming that only the neurons tuned to a given stimulus feature contribute to its encoding. Recent experimental work questioned this assumption, showing that neurons with no apparent tuning can sometimes contribute to information coding. However, key questions remain unanswered. First, how do the untuned neurons contribute to information coding, and could this effect rely on those neurons having weak tuning that was overlooked? Second, does the function of a neural circuit ever benefit from having some neurons untuned? Or should every neuron be tuned (even weakly) to every stimulus feature? Here, I use mathematical calculations and analyses of data from the mouse visual cortex to answer those questions. First, I show how (and why) correlations between neurons enable the untuned neurons to contribute to information coding. Second, I show that neural populations can often do a better job of encoding a given stimulus feature when some of the neurons are untuned for that stimulus feature. Thus, it may be best for the brain to *segregate* its tuning, leaving some neurons untuned for each stimulus feature. Along with helping to explain how the brain processes external stimuli, this work has strong implications for attempts to decode brain signals, to control brain-machine interfaces: better performance could be obtained if the activities of all neurons are decoded, as opposed to only those with strong tuning.

## Introduction

When you look at a picture, signals from your eyes travel along the optic nerve to your brain, where they evoke activity in neurons in the thalamus and visual cortex. As sensory systems neuroscientists, we ask how these patterns of stimulus-evoked brain activity reflect the outside world – in this case, the picture at which you are looking. Other related work asks how patterns of activity in different parts of the brain reflect motor commands sent to the muscles. Answers to these questions are important both for basic science, and for brain-machine interface technologies that either decode brain activity to control prosthetic limbs or other devices [1, 2, 3], or stimulate the brain to alleviate sensory deficits [4, 5].

For decades, researchers have addressed these information coding questions by recording neural activity patterns in animals while they are being presented with different stimuli, or performing different motor tasks. That work revealed that many neurons in the relevant brain areas show firing rates that depend systematically on specific features of the stimulus presented to the individual, or on the motor task (e.g., motion direction of a moving visual stimulus). This neural “tuning” underlies the ability of these neural circuits to encode information about the stimulus and/or behavior. At the same time, many neurons appear to be untuned for each stimulus feature, thus showing little or no systematic variation in their firing rates as that stimulus (or behavioral) feature is changed [6]. (Note that neurons that are untuned for one stimulus feature could be tuned to a different one). These untuned neurons are typically ignored in studies of neural information coding because it is presumed that neurons do not contribute to encoding stimulus features to which they have no tuning [7]. Instead, data collection and analysis are typically restricted to those neurons that are tuned to the stimulus feature(s) of interest to the experimenters (for example, consider the selection criteria used by [8, 9]).

Recently, researchers have begun to question that assumption: analyses of neural data in the prefrontal cortex [10], somatosensory cortex [11], and auditory cortex (Insanally et al., 2017 cosyne abstract), show that even neurons with no obvious tuning to a given stimulus feature can nevertheless contribute to the population code for that stimulus feature. These findings are intriguing, because they suggest that – by virtue of our ignoring the untuned neurons – our understanding of neural population coding might be incomplete. One of the prior studies also provides nice intuition, based on pairs of neurons, for why untuned neurons could matter for population coding [10].

At the same time, several deep questions remain unanswered: First, are the impacts of these putatively untuned neurons on population coding due to weak tuning that is nevertheless below the threshold the experimenters set for calling neurons tuned (vs untuned)? Second, does the intuition from [10] for why untuned neurons could improve population coding, based on pairs of neurons, hold for larger population sizes and for different correlation structures? Given prior work showing that population coding intuition can often fail to extrapolate from pairs of neurons to larger populations [12], this is a potentially important question. Third, can mixed populations of tuned and untuned neurons have a functional advantage over populations containing only tuned neurons?

To answer these questions, I used theoretical calculations, and then verified the predictions from those calculations by analyzing 2-photon imaging data collected in the visual cortices of mice that were shown drifting grating stimuli [13]. For the theoretical calculations, I used a common mathematical model of the neural population responses to sensory stimulation, known as “tuning curve plus noise” [14, 15, 16, 17, 12, 18, 19, 9, 20, 21, 22, 19, 23].(Note that, depending on the structure of that noise, some of these models show information that saturates with increasing population size, while others do not.) This class of model describes key features of sensory neural responses: the stimulus tuning (or lack thereof) of individual neurons; the trial-by-trial deviations (or “noise”) in the neural responses [9, 24, 25, 26, 27, 28]; and the potential for that noise to be correlated between neurons [29, 30, 9, 31, 32, 29, 33, 34, 35, 36, 37, 38]. For different conditions – for example, including vs. excluding untuned neurons – I computed the amount of information about the stimulus that is encoded in the population firing patterns. By comparing the information across conditions, I characterized the impact that untuned neurons can have on the neural population code.

Because the untuned neurons in the theoretical model really have no stimulus tuning, these calculations enabled me to demonstrate conclusively that strictly untuned neurons really can contribute to population coding: I show that this effect holds for a wide range of correlation structures, and neural population sizes. Moreover, by studying the information coding of neural populations containing different fractions of tuned vs untuned neurons, I demonstrated that mixed populations can encode stimulus information better than populations containing only tuned neurons. Finally, I used decoding analyses applied to data collected in the visual cortices of awake mice to validate the key predictions of the theory: excluding putatively untuned neurons hinders decoding; and decoding random groups of neurons (both tuned and untuned) yields better performance than does decoding neural populations of the same size, but containing only tuned neurons.

## Results

I first study a theoretical model of information coding in neural populations, to understand whether and how untuned neurons contribute to information coding. I then validate the main predictions from the theory by analyzing data collected in mouse visual cortex.

## Theoretical analysis

### The role of untuned neurons in sensory information coding

To investigate the role of untuned neurons in sensory information coding, I studied populations of neurons that encode information about the motion direction of a visual stimulus via their randomly shaped and located tuning curves (Fig. 1A). Many different population sizes were considered. For each population, 70% of the neurons were tuned, and the other 30% were untuned. (These numbers match the fraction of well-tuned neurons selected for analysis in a recent population imaging study [38], and are comparable to the fraction of tuned neurons in the experimental data that I analyzed. I later consider populations with different fractions of untuned neurons.) The untuned neurons had flat tuning curves that did not depend on the stimulus – see the dashed lines in Fig. 1A. The neurons had Poisson-like variability: for each cell, the variance over repeats of a given stimulus was equal to the mean response to that stimulus. This mimics the experimentally observed relation between means and variances of neural activities [25, 23]. The variability was correlated between cells, and the correlation coefficients were chosen to follow the “limited-range” structure reported experimentally [30, 18, 39, 40, 41], and used in previous theoretical studies [14, 15, 16, 20]. With this structure, the correlation coefficients were large for neurons with similar preferred directions, and smaller for neurons with very different preferred directions (see Methods and Fig. 1B). (I later consider models with different correlations structures – that lead to information that saturates with increasing population size.) I did not consider negative noise correlations here – even though those correlations structures can in some circumstances have beneficial effects on population coding [15, 16] – largely because experimentally observed noise correlations are typically positive.

**Figure 1:**
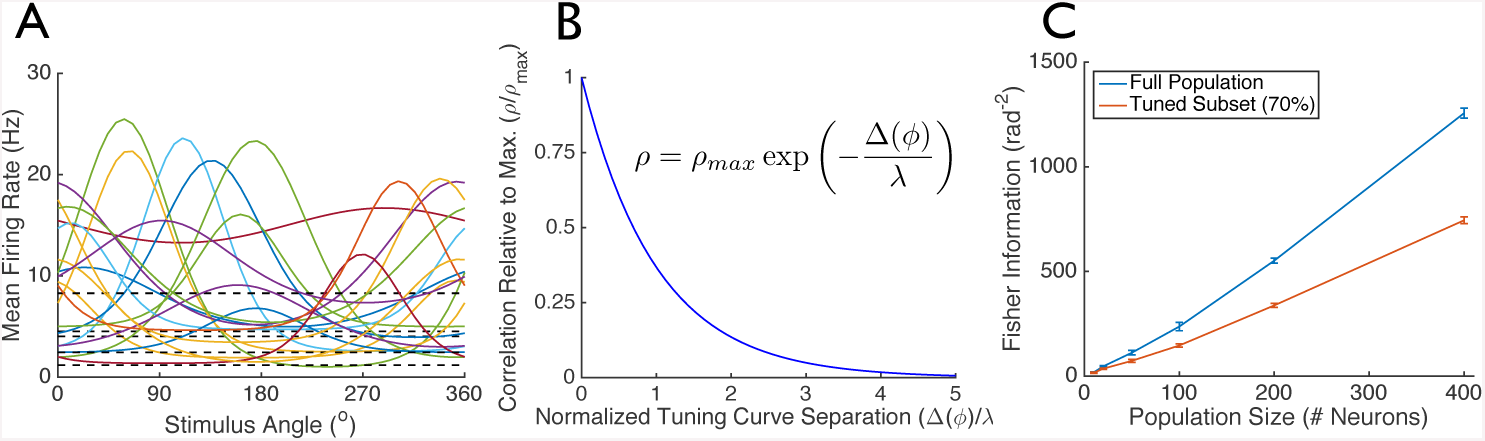
Untuned neurons can play an important role in sensory information coding. (A) I considered populations of neurons with randomly shaped and located tuning curves. Of those neurons, 70% were tuned to the stimulus, whereas 30% were untuned – their mean firing rates do not depend on the stimulus (dashed black lines in panel A). The neurons’ trial to trial variability was Poisson-like and correlated between neurons. (B) These correlations followed the “limited-range” structure with *ρ*_*max*_ = 0.75 and *λ* = 0.5 radians (29^°^). The mean correlation coefficients (averaged over neurons) were 0.12, which is comparable to values reported in primary visual cortex [30]. (Modifying these values did not qualitatively change the results – see Fig. S1). (C) For different sized populations, I computed the Fisher information, which quantifies how well the stimulus can be estimated from the neural population activities. The different lines correspond to: the Fisher information for the full neural populations (blue); and the Fisher information for the tuned 70% of the populations (red). Data points are mean ± S.E.M., computed over 5 different random draws of the tuning curves.

For each population, I computed the Fisher information (Fig. 1C, blue curve), which quantifies how well an observer – like a downstream neural circuit – can estimate the stimulus direction angle from the neural activities (see Methods). I compared that with the Fisher information obtained from only the tuned subset of neurons – in other words, the information that would be obtained if the untuned cells were ignored (Fig. 1C, red curve). The difference was stark. Ignoring the untuned neurons leads to a dramatic underestimate of the encoded stimulus information. This emphasizes that, despite their lack of direction dependence, the untuned neurons can still contribute significantly to the population code for direction.

Because the correlation coefficients in Fig. 1 did not depend on the stimulus, it is not the case that the untuned neurons themselves encode information indirectly, through their second-order statistics (as was the case in the theoretical model of [42]). This point is emphasized in Fig. 4, where the information in the population goes to zero as the fraction of untuned neurons approaches 100%. This suggests the question of how untuned neurons contribute to neural information coding. While the untuned neurons’ activities do not reflect the stimulus, they do reflect the trial-specific noise in the tuned neurons’ activities (because they are correlated). Accordingly, a downstream readout – like the circuit receiving these neural spikes – can obtain a less noisy estimate of the stimulus by using the untuned neurons’ activities to estimate the noise in the activities of the tuned neurons, and subtracting that noise estimate from the observed firing rates. Ignoring untuned neurons leads to the loss of the information available through this “de-noising”.

To illustrate this point, I considered a pair of neurons, one of which is tuned to the stimulus direction (Fig. 2A); this geometrical argument is also made by [10]. In response to stimulation, the neurons give noisy responses, and that noise is correlated between the two cells. When plotted in the space of the two cells’ firing rates, the distributions of neural responses to each stimulus direction are defined by ellipses, shown in Fig. 2B. (These are the 1 standard deviation probability contours.) The correlation between cells is reflected in the fact that these ellipses are diagonally oriented. These ellipses are relatively disjoint, meaning that the neural responses to the different stimuli have little overlap, and so it is relatively unambiguous to infer from the neural firing rates which stimulus was presented. For contrast, consider the neural activities observed when the untuned neuron is ignored. In that case, only the tuned neuron is observed, and the distributions in its responses to the different stimuli overlap substantially (Fig. 2B, right vertical axis). This means that, based on only observations of the tuned cell, the stimulus cannot reliably be determined. Ignoring the untuned neuron leads to a loss of stimulus information.

**Figure 2:**
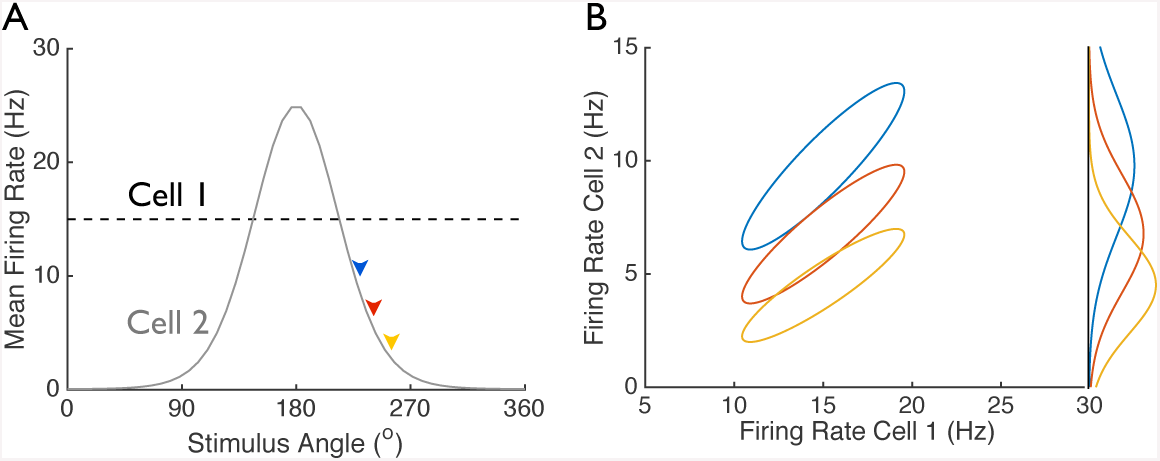
Untuned neurons can shape noise, improving the population code. (A) Two neurons’ tuning curves are shown; cell 1 is untuned. In response to stimulation, the cells give noisy responses. That noise is correlated between the two neurons, with a correlation coefficient of 0.9. (B) The distribution of noisy responses to each stimulus is described by an ellipse in the space of the two neurons’ firing rates. The stimulus values are indicated by arrows in panel (A). The ellipses are well separated, meaning that the stimuli can be readily discriminated based on the two cells’ firing rates. If the untuned cell is ignored, then only the tuned cell is observed. The distribution of the tuned cell’s firing rate in response to each stimulus is shown along the right vertical of panel (B). Because those distributions overlap substantially, the stimulus cannot be readily discriminated based only on the firing rate of the tuned cell. This geometrical argument is also made by [10].

Because the untuned neurons’ contribution to the population code relies on their activities reflecting the single-trial noise in the activities of the tuned cells, the untuned neurons do not contribute to population coding if they are independent of the tuned neurons [10]. To demonstrate this point, I repeated the analysis from Fig. 1 (above), but made the untuned neurons uncorrelated from each other and from the tuned neurons. In that case, the untuned neurons do not contribute to the population code: the full population and the tuned subset both have the same amount of stimulus information (Fig. 3A).

This contribution of untuned neurons to the population code can be understood via the cartoon in Fig. 3B (lower), which shows the distribution of population responses to 3 different stimuli. In the cartoon, cell 1 is untuned, whereas the rest of the cells are tuned. This means that, as the stimulus changes, the mean responses change along the plane orthogonal to the cell 1 axis. Because the untuned neuron is correlated with the tuned ones, the noise distributions are tilted along the vertical axis. In this configuration, the distributions do not overlap very much. If, however, the untuned neuron is made independent from the tuned ones (as in Fig. 3A), the vertical tilt goes away, causing much more overlap in the response distributions. In other words, when the untuned neurons are correlated with the tuned ones, they improve the population code by separating the responses to different stimuli. This effect disappears when the untuned neurons are independent of the tuned ones. Similar geometrical arguments are made by [10].

**Figure 3:**
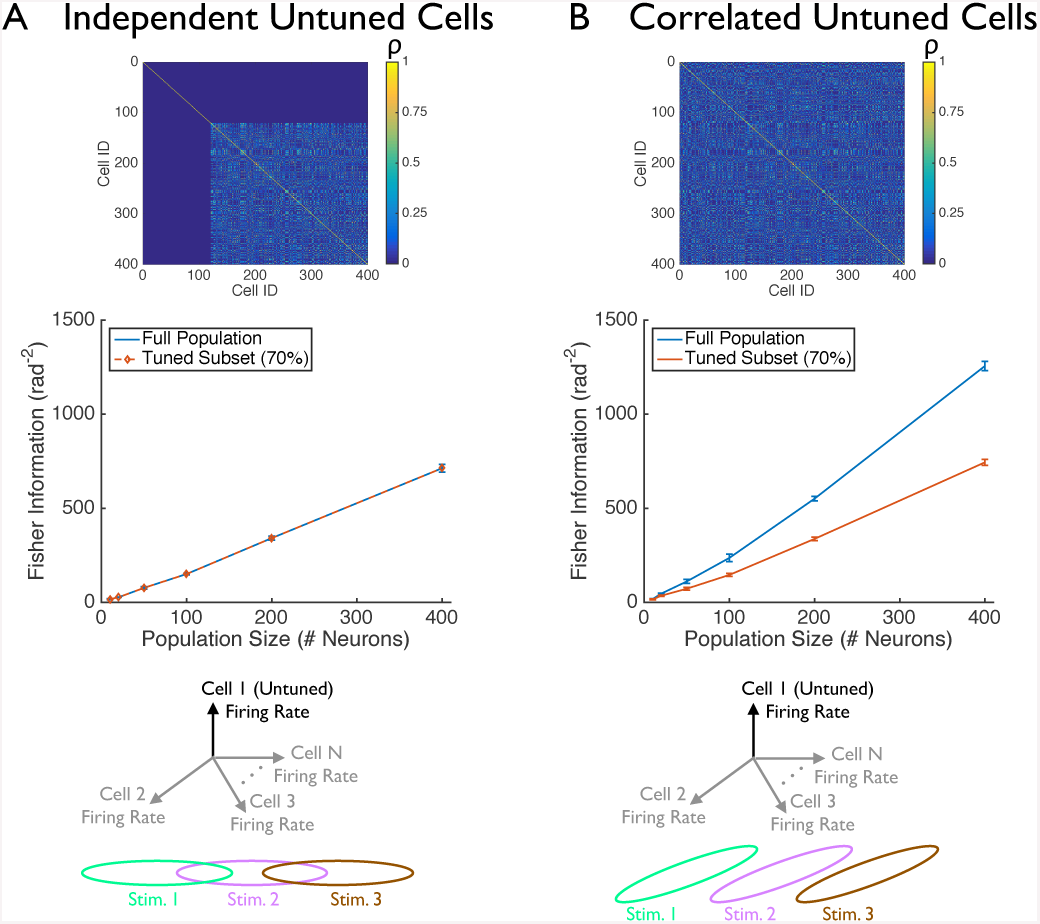
Untuned neurons improve population coding when they are correlated with the tuned neurons. (A) I repeated the analysis from Fig. 1C, and modified it so that the untuned neurons were independent of each other and of the tuned neurons. (B) For comparison, I also show again the results from Fig. 1C. As in Fig. 1, 70% of the neurons in each population were tuned to the stimulus, and 30% were untuned. Upper panels show correlation matrices for 400-cell populations: cells 1 through 120 are untuned, whereas the remainder were tuned. Center panels show the Fisher information for the full neural populations (blue), or for the tuned subsets of neurons in each population (red). (Data points shown are mean ± S.E.M., computed over 5 different random draws of the tuning curves). The cartoons in the lower panels illustrate why these two different correlation structures lead to untuned neurons having such different effects on the population code (see text, and [10]). The cartoons show the space of neural firing patterns: each axis is the firing rate of a different neuron. The vertical axis is the firing rate of an untuned neuron. The other axes are the firing rates of tuned cells. Ellipses represent the 1 standard deviation probability contours of the neural population responses to the 3 different stimuli.

The results in Figs. 1 and 3 are for neural populations in which information increases without bound as the population size increases. However, in many large neural populations, information saturates with increasing population size [19]. Accordingly, it is important to check that the same results hold for these population codes. As shown in Fig. S2, populations with *information-limiting* correlations, show the same main effect as the populations with limited-range correlations studied in Figs. 1 and 3: ignoring untuned neurons leads to a reduction in information.

### Mixed populations of tuned and untuned neurons can encode stimulus information more effectively than populations containing only tuned neurons

In the preceding analyses, I showed that neurons with no stimulus tuning can contribute to the population code: ignoring them entails a loss of stimulus information. Here, I turn to the question of why the brain might contain those neurons at all. In other words, is there a functional benefit to including untuned neurons in a population vs having only tuned neurons?

To answer this question, I repeated the analysis from Fig. 1 – again, using populations of heterogeneously tuned neurons with limited-range correlations – but altered the fraction of untuned neurons in each population. The maximum information values were obtained with around 30% of neurons being untuned; this effect was larger in larger populations (Fig. 4A). Because the maximum information does not occur when all of the neurons are tuned (corresponding to an untuned neuron fraction of 0), this analysis shows that neural populations can be made more informative by replacing tuned neurons with untuned ones. Note that the fraction of untuned neurons that maximizes information depends on the structure of the correlations in the population. For example, with no correlations, maximum information is obtained when all neurons are tuned.

**Figure 4:**
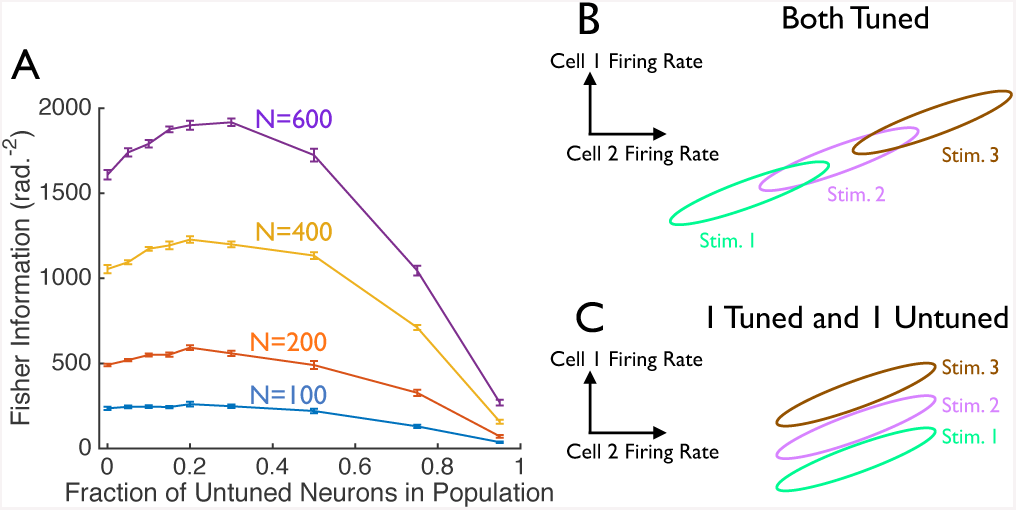
Populations containing some untuned neurons can encode more information than ones with only tuned neurons. (A) I repeated the calculations from Fig. 1, but with different fractions of untuned neurons in each population. For several different population sizes (indicated on the plot), the Fisher information is shown as a function of the fraction of untuned neurons in the population. As in Fig. 1, each population had limited-range correlations, with average correlation coefficients that are in the physiological range. Error bars are the S.E.M. over 5 random sets of different tuning curves. (B and C) Cartoon showing how mixed populations containing tuned and untuned neurons can be better at encoding information than populations containing only tuned neurons. In response to 3 different stimuli, I show the 1 standard-deviation probability contours in the responses of a pair of neurons. In all cases, the neurons have Poisson variability, and correlated noise. In panel (B), both cells are tuned to the stimulus, whereas in panel (C), cell 1 is tuned to the stimulus, and cell 2 is untuned.

How can mixed populations of tuned and untuned neurons be better at encoding information than populations of the same size but containing only tuned cells? The cartoon in Figs. 4B and C provides some intuition. In both cases, the distributions of firing rates of two neurons are shown, in response to 3 different stimuli (similar to Fig. 2B): ellipses indicate 1 standard-deviation probability contours. In Fig. 4B, both cells are tuned to the stimulus, and the centers of the ellipses are correspondingly displaced relative to each other along both the vertical, and the horizontal, axes of the plot. With this geometrical configuration, the ellipses corresponding to different stimulus-evoked responses overlap substantially: that overlap means that there is ambiguity in determining the stimulus from the neural responses, and so the population code has relatively low information. Fig. 4C differs from Fig. 4B only in the tuning of cell 2: in Fig. 4C, cell 2 is untuned, whereas in Fig. 4B, it was tuned. This means that, in Fig. 4C (where only one of the cells is tuned to the stimulus), the different stimulus-evoked response distributions are displaced relative to each other only in the vertical direction, and not the horizontal one. Owing to the diagonal orientation of the ellipses, there is less overlap between the different response distributions in Fig. 4C than 4B. Consequently, the pair of neurons in Fig. 4C (one of which is untuned) is better at encoding stimulus information than the pair of neurons in Fig. 4B (both of which are tuned to the stimulus).

The cartoon in Figs. 4BC shows how the presence of untuned neurons can improve the population code: including untuned neurons modifies the *signal correlation* structure (the correlation between neurons in the stimulus-evoked mean responses) relative to the case where both neurons are tuned. And because the relationship between the signal and noise correlations determines the population coding efficacy [12, 17, 15], this modification can improve the population code overall.

Under what conditions do mixed populations containing tuned and untuned neurons encode stimulus information better than populations containing only tuned cells? To answer this question, I performed mathematical analyses – described in detail in the Methods – that identify conditions where the population code can be made more informative by replacing a tuned neuron (neuron *k*) by an untuned one. Those analyses showed that making neuron *k* untuned will improve the population code whenever the following inequality holds:

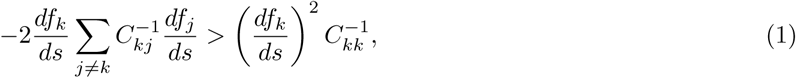

where 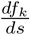 is the slope of the tuning curve of neuron *k*, and *C* is the covariance matrix of the neural variability. Intuitively, this equation compares the loss of information from removing the tuning of neuron *k* (the right-hand side of Eq. 1), with the gain in information from the noise-shaping effect shown in Figs. 4BC (the left-hand side of Eq. 1). Whenever the gain exceeds the loss, it is beneficial to make neuron *k* untuned. Note that the inequality in Eq. 1 will not necessarily be satisfied by all sets of neural tuning curves and covariance matrices. Consequently, it is not guaranteed that including untuned neurons will *always* improve the population code. However, under the condition specified by Eq. 1, there is a functional benefit to including untuned neurons in a population.

### Analysis of *in vivo* neural activities

The theoretical work described above makes a key prediction: the ability to decode a stimulus from the evoked neural population activities could be improved if untuned neurons are included in those populations, as opposed to being ignored. To test this prediction, I analyzed data from 2-photon Ca^2^+ imaging recordings done in primary visual cortex of awake mice (data from [13]) whose neurons expressed the genetically encoded calcium indicator GCaMP6f. The mice were presented with stimuli consisting of gratings drifting in 8 different directions, and the fluorescence levels of 𝒪(100) neurons were observed in each experiment. I analyzed the data from 46 such experiments.

For each stimulus presentation and neuron, I extracted the mean fluorescence change during the stimulus presentation, relative to the fluorescence in the period before the stimulus presentation: this Δ*F/F* value measures the stimulus-induced change in neural activity. I then computed the neurons’ tuning curves by averaging these Δ*F/F* values over all trials in which the stimulus drifted in each direction. Some of the neurons had well-defined direction tuning curves (Fig. 5A), whereas others were relatively untuned (Fig. 5B). Following [43, 9], I categorized these cells as tuned or putatively untuned (hereafter referred to simply as *untuned*) based on their direction selectivity indices (see Methods). Between the 46 experiments, 5379/8943 ≈ 60% of the neurons were classified as being tuned for direction. (In the population coding analyses discussed below, I consider several different thresholds for labelling neurons as “tuned” vs “untuned”.)

**Figure 5:**
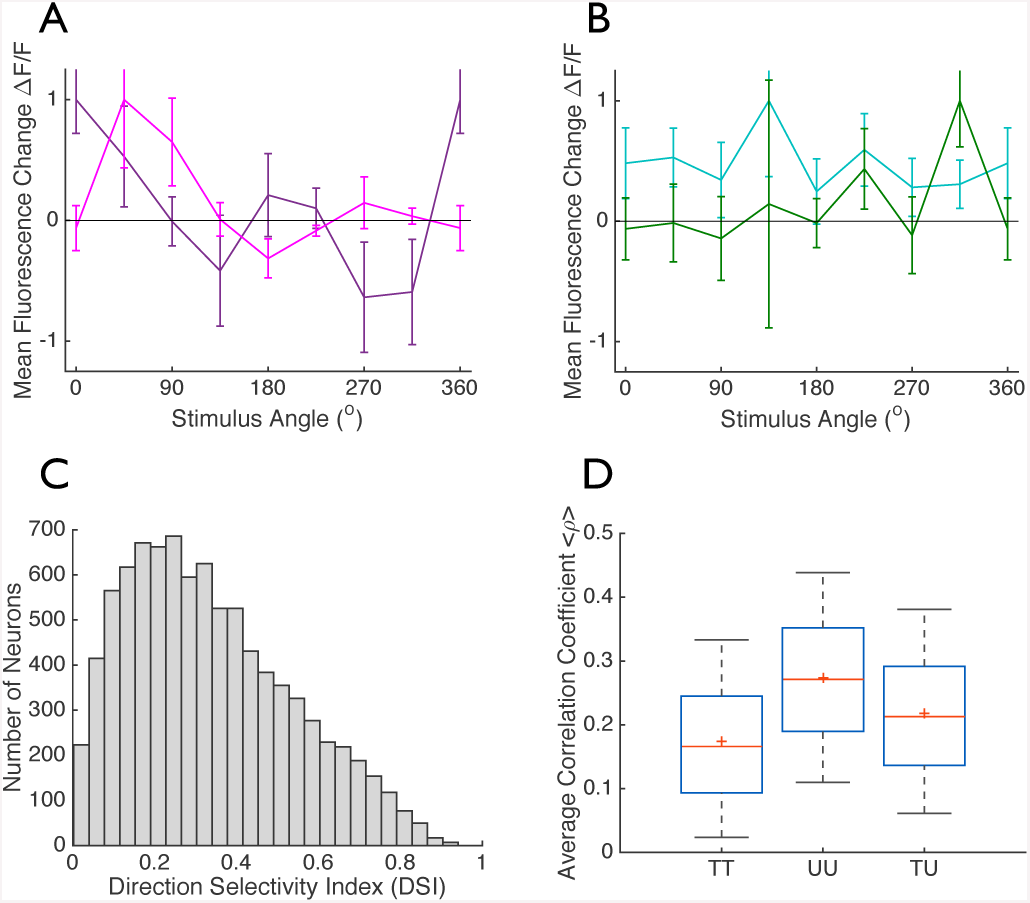
Tuned and untuned neurons are correlated *in vivo*. Neurons’ responses to drifting grating stimuli were measured using 2-photon Ca^2+^ imaging. (A) Tuning curves for two direction tuned neurons. (B) Tuning curves of two untuned neurons. Markers show mean Δ*F/F* ± S.E.M, calculated over 75 trials of each stimulus direction. (C) Direction selectivity indices for the 8493 neurons whose stimulus-evoked responses were measured. (D) The distributions of correlation coefficients for cell pairs of different types: where both cells were direction tuned (“TT”; *n* = 391833 pairs); where both cells were untuned (“UU”; *n* = 150801 pairs); and where one cell was tuned and one was untuned (“TU”; *n* = 434752 pairs). Each box plot shows the median, the range (maximum and minimum indicated by black bars), and the boundaries of the 25^*th*^ and 75^*th*^ percentiles (blue box) of the distributions.

Along with the tuning, I measured the correlations in the cells’ trial-to-trial variability over repeats of each stimulus. These “noise correlations” are shown for all pairs of simultaneously observed neurons (Fig. 5D). The correlation coefficients were typically positive for pairs of tuned neurons (“TT”), pairs of untuned neurons (“UU”), and mixed pairs consisting of one tuned and one untuned neuron (“TU”). Because there were correlations between the tuned and untuned neurons, the theory predicts that stimulus decoding could be improved by including the untuned neurons, as opposed to ignoring them.

To test this prediction, I used the logistic regression method of [44] to take in vectors of neural activity recorded in response to one of 2 different stimuli, and to identify the stimulus from those neural responses. The classifier was trained on 75% of the data, and I tested the performance on the remaining held-out 25% of the data. (see Methods for details). To test the performance, I computed the fraction of trials in the test dataset on which the stimulus was correctly identified.

I first performed this analysis on the full populations of recorded neurons – including the tuned and untuned ones – and compared this to the decoding performance when only tuned neurons were used by the decoder. Using all of the neurons resulted in 10 ± 1% (mean ± S.E.M) better decoding performance (*p* = 8.9 ×; 10^−11^, paired one-sided t-test; and *p <* 10^−8^, non-parametric binomial test of significance) than did using only the tuned neurons (Fig. 6A).

**Figure 6:**
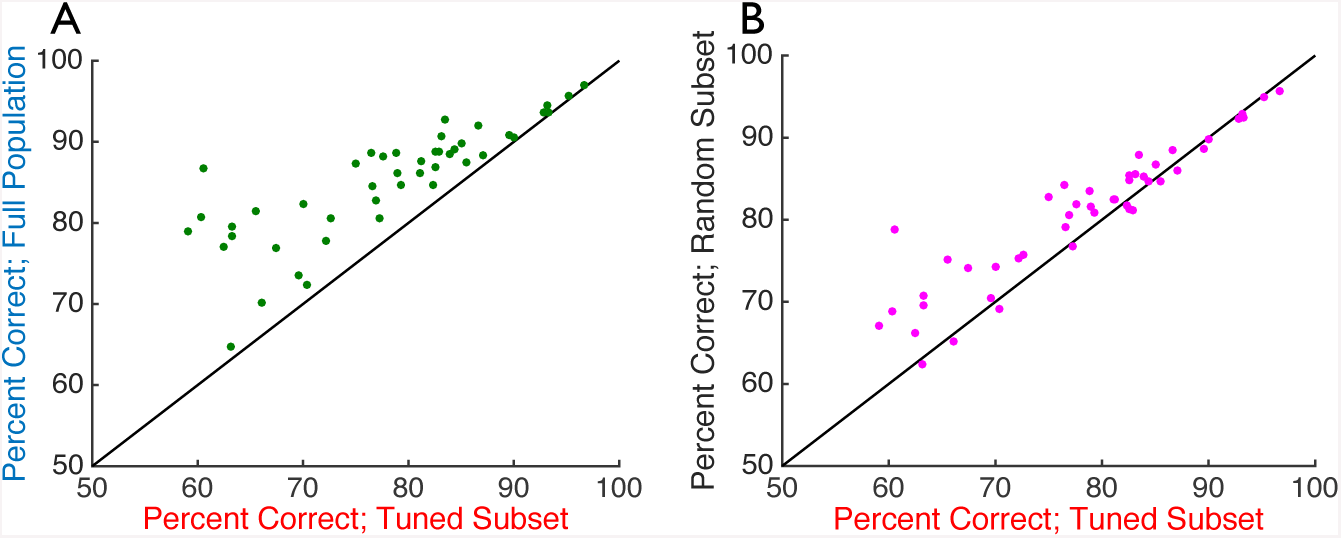
Untuned neurons can enhance information coding in *in vivo* neural populations. I used logistic regression to perform pairwise discrimination on the population response vectors, to determine which of 2 different stimuli caused each response. I repeated this analysis for all possible pairs of stimuli: reported values are the percentage of trials for which the stimulus was correctly identified, averaged over all possible pairings (there is one data point per experiment). (A) Decoding accuracy when the full population response vectors were decoded (vertical axis) vs. when only the tuned subsets of the neurons are seen by the decoder (horizontal axis). (B) Decoding accuracy when random subsets of the neurons (of the same size as the tuned subset, but containing both tuned and untuned neurons) are input to the decoder (vertical axis) vs. when only the tuned subsets of the neurons are seen by the decoder (horizontal axis). Chance performance for this binary discrimination task is 50%. Diagonal line denotes equality.

Next, I asked whether – as in the theoretical calculations in Fig. 4 – populations that include both tuned and untuned neurons could yield better decoding vs populations of the same size but containing only tuned cells. To answer this question, I extracted a random subset of the neurons from each population, that was the same size as the set of tuned neurons within that population. I then performed the logistic regression analysis on these random subsets, and compared the performance with that which was obtained on the tuned subsets (Fig. 6B). On average, the decoding performance was 4 ± 1% (mean ± S.E.M.) better using the random subsets vs the fully tuned ones, a modest but statistically significant difference (*p* = 1.7 ×; 10^−5^, paired single-sided t-test; and *p* = 0.027, non-parametric binomial test of significance). While this effect was modest in size, the sign was surprising: prior to performing this study, I would not have anticipated that populations containing untuned neurons could form better population codes than do populations the same size but containing only tuned cells.

It is important to check that the results in Fig. 6 do not depend on the specific criterion used to distinguish tuned from putatively untuned neurons. Consequently, I repeated the analysis from Fig. 6 with several different criteria (see Methods and Figs. S4-S5). These results are all in qualitative agreement with those of Fig. 6: regardless of the specific criterion that is used, the putatively untuned neurons contribute to the population code, and mixed populations of tuned and untuned neurons encode more information than do populations of the same size but containing only tuned neurons. Notably, the sizes of these effects decrease as one uses progressively less strict selection criteria. This is because, with less strict selection criteria, fewer putatively untuned neurons are excluded. In the limit where all neurons are labelled as tuned, there are no untuned neurons, and thus they have no effect.

## Discussion

I showed that, when the variability in neural responses to stimulation is correlated between cells, even neurons whose firing rates do not depend on the stimulus (“untuned” neurons) can contribute to sensory information coding: unlike prior work [10], I showed that this is true for a range of population sizes and correlation structures. Moreover, in at least some cases, populations with both tuned and untuned neurons can convey more information about the stimulus than do populations of the same size but containing only tuned neurons. These effects were observed in both a theoretical model (Figs. 1-4), in and in large population recordings from mouse visual cortex (Fig. 6). These experimental findings were not sensitive to the specific criterion used to define neurons as being tuned vs untuned (Figs. S4-S5).

These results have two main implications. First, our understanding of how the sensory systems encode information about the outside world is likely to be incomplete unless it includes the contributions of all neurons, regardless of whether or not they appear to be tuned to the stimulus feature of interest. This means that current practices, in which putatively untuned neurons are ignored during data collection and analysis, might be hindering progress. Moreover, because there is not always a clear distinction between tuned and untuned neurons (Fig. 5C: histogram is unimodal) – and this effect is confounded by noise in the experimental measurements – selection criteria are largely arbitrary. This experimental noise also means that is is nearly impossible to know whether there really are strictly “untuned” (as opposed to only weakly tuned) neurons in the brain. Either way, the results shown here suggest that, rather than discarding neurons with weak (or zero) tuning, it is better to simply include all the neurons in the analysis: even in the extreme case, where those neurons really have *no* tuning, they can still contribute to the population code. This last point applies especially to decoding population activities to control brain-machine interface devices: better performance could be obtained by decoding all neurons, as opposed to decoding only the well-tuned ones (Fig. 6A). At the same time, decoding large neural populations using limited amounts of training data can be subject to overfitting; in that case, ignoring the untuned neurons can effectively reduce the amount of data needed to train the decoder. This could potentially lead to improved decoding performance. However, it is important to note that there are other forms of regularization (like Ridge, or Lasso), that can help overcome this challenge, without requiring one to *a priori* select neurons to ignore. Based on the results of this study, my recommendation is to use regularizers like Ridge or Lasso, but include all recorded neurons in the decoding.

Second, because adding untuned neurons can increase the stimulus information (Figs. 4, 6), there might be a functional benefit to having some neurons in a population that are untuned for each stimulus feature. In other words, rather than having every neuron be somewhat tuned (even weakly) to every stimulus feature, it may be better for the tuning to be *segregated*, such that sub-populations remain untuned for each stimulus feature. This is related to previous observations that heterogeneous tuning curves could confer advantages on the population code [45, 16]. Those previous studies did not, however, consider the role of untuned neurons in the population code. It is important to note, however, that no brain area can encode more stimulus information than it received from its inputs [46, 23]. This is the *data-processing inequality*, and it implies that there is not a limitless increase in information to be obtained by adding large numbers of untuned neurons to neural circuits.

While I derived these results in a theoretical (and experimental) setting with only one stimulus feature being encoded, the mathematical results generalize to larger numbers of different stimulus features. Notably, the Fisher information (Eq. 2) can be computed for any given stimulus feature, even if there is more than one of interest. This means that our results can be applied to each stimulus feature, one at a time, and thus generalize directly to the problem of encoding multiple stimulus features.

Observations related to those presented here have also been made by Insanally and colleagues (Cosyne 2017 abstract), and by [10, 11] based on analyses of *in vivo* neural data: those observations were in the auditory system, somatosensory system, and prefrontal cortex. There, as in the analysis of visual cortical data presented here, it is hard to distinguish weakly tuned neurons from purely untuned ones, and thus difficult to isolate the coding benefits of putatively untuned neurons due to noise shaping, vs those due to non-zero tuning, that is nonetheless under the chosen threshold. (However, the fact that the results are not sensitive to the specific criterion uses to label neurons as tuned vs untuned does help to make this distinction: Figs. S4, S5). This complication highlights the value of the theoretical work presented here (Figs. 1-4): in the model, the untuned neurons really have no stimulus dependence, enabling us to shown that even neurons with *no* tuning can contribute to sensory information coding.

For large neural populations, an astronomically large number of different correlation patterns are possible (and this problem is confounded when one includes correlations of higher order than the pairwise ones considered here [47, 48]). Accordingly, it was not possible to simulate all possible correlation patterns in the theoretical study. Thus, it is natural to ask how general the results are over different correlation structures. Here, the fact that I saw consistent effects in the experimental data (Fig. 6) as in the theoretical model with limited range correlations (Fig. 1), and the model with information-limiting correlations (Figs. S2, S3), argues for the generality and applicability of the findings.

The experimental data studied here (Figs. 5 and 6) were recorded while mice were passively viewing the visual stimulus. Consequently, the correlated fluctuations in visual cortical neural activity could correspond to changes in attention, arousal, or other factors that were not controlled in the experiment. As a result, it is important for future work to assess the role of untuned neurons under experimental conditions with more carefully controlled behavior; that work is beyond the scope of this study.

Adding neurons to a population can never decrease the amount of encoded stimulus information: because a downstream read-out could always choose to ignore the added cells, those cells can at worst contribute zero information. Consequently, untuned neurons can never *hinder* the population code. This means that the potential effects of untuned neurons on population coding range between no contribution (Fig. 3A), and positive contributions at least as large as those seen in Fig. 1C (i.e., at least 70% increase in information available by including vs. ignoring untuned neurons). There may be other cases in which the positive contributions of untuned neurons are even larger.

It is important not to interpret the results presented here as implying that neural tuning is not essential to sensory information coding. Whereas the theoretical model of [42] can encode stimulus information via changes in the correlations between neurons, that effect is not responsible for the results shown here. Notably, for the theoretical calculations in Figs. 1-4, the correlations do not depend on the stimulus, yet nevertheless the untuned neurons contribute to the population code. Underscoring this point is the fact that, if there are no tuned neurons in our models, there is no stimulus information (Fig. 4: information approaches zero as the fraction of untuned neurons approaches 1).

I conclude by noting that, even when untuned neurons do not by themselves encode information about the stimulus, they can shape the noise in the population responses, thereby improving the population code overall. Thus, untuned neurons are not irrelevant for sensory information coding.

## Methods

I first discuss the theoretical calculations, and then the analysis of experimental data.

## Theoretical Calculations

### Modeling the stimulus-evoked neural responses, and the information encoded

I considered for simplicity a 1-dimensional stimulus *s* (for example, the direction of motion of a drifting grating). In response to the stimulus presentation, the neural population displays firing rates 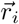, where the index *i* denotes the trial. (Each element of the vector 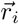 is the firing rate of a single neuron). These responses have two components. The first, 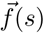, is the mean (trial-averaged) response to stimulus *s*, whereas the second component, 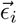, represents the trial-by-trial fluctuations, or “noise” in the neural firing rates.

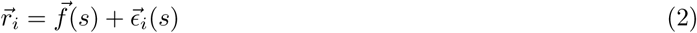

Following [16, 12, 23], the tuning curves were generated using Von Mises distributions [16],

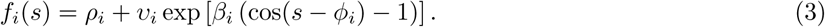

For each tuned cell, the amplitudes, *v*, widths, *β*, peak locations, *ϕ*, and baseline offsets, *ρ*, were drawn independently from uniform distributions with the following ranges,

- *v*: 5–51
- *β*: 1–6
- *ϕ*: 0–2*π*
- *ρ*: 0–1

The untuned neurons had *β* values of zero; and their other tuning curve parameters were drawn from the same distributions as were those of the tuned cells (above).

The neurons’ noise variances were chosen to match the mean responses, in accordance with experimental observations of Poisson-like variability. I considered different patterns of inter-neural correlation, as described below.

For each set of tuning curves and correlations, I used the typical linear Fisher information measure, *I(s*), to quantify the ability of downstream circuits to determine the stimulus, *s*, from the noisy neural responses on each trial 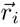 [14, 15, 16, 17, 12, 18, 20, 22, 19, 21, 23]:

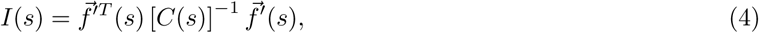

where the prime denotes a derivative with respect to the stimulus, the supserscript *T* denotes the transpose operation, and 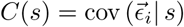 is the covariance matrix of the noise in the neural responses to stimulus *s*. For all calculations, I checked that the correlation (and covariance) matrices were positive semi-definite (thus being physically realizable) before performing the Fisher information calculations.

To compute the information for a subset of a neural population, I extracted the block of the covariance matrix, and the elements of the vector 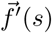, that correspond to the neurons in that subset. I then used those values in Eq. 2.

For all of the information values presented here, I computed the information for each of 50 different stimulus values, evenly spaced over [0°, 360°]. The reported values are averages over these 50 stimuli. This accounts for the fact that Fisher information *I(s*) is a *local* quantity which varies from stimulus to stimulus. By averaging over many stimuli, I avoid the possibility that the reported information values might be atypical, and affected by the specific stimulus at which the information was calculated.

### Limited-range correlations

The elements of covariance matrix *C(s*) were 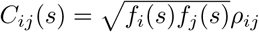, where *ρ*_*ij*_ is the correlation between cells i and j. The factor of 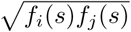 ensures that the neurons have Poisson variability (variance of noise is equal to mean firing rate, meaning that standard deviation of noise is equal to square root of mean firing rate).

The correlation coefficients *ρ*_*ij*_ were calculated as (Fig. 1B)

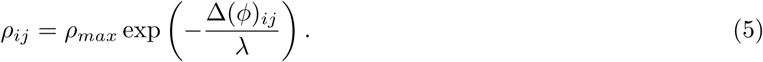

The tuning curve separation for each cell pair was computed as Δ(*ϕ*)_*ij*_ = *|* arccos [cos(*ϕ*_*i*_ *-ϕ*_*j*_)] *|*, where *ϕ*_*i*_ and *ϕ*_*j*_ are the cells’ preferred direction angles (the locations of their tuning curve peaks). This formula accounts for the fact that angles “wrap” around the circle: so values of 10° and 350° have a separation of 20° (and not 340°).

For the untuned neurons, their preferred stimulus angles were randomly assigned, uniformly over the range [0°, 360°].

### When do untuned neurons improve population coding?

Here, I derive Eq. 1 from the main text, which specifies the conditions under which including untuned neurons in a population improves its ability to encode stimulus information. To do this, I compute the information in the neural population, and the information that would be obtained if one of the neurons were to be made untuned. I then ask when the information increases as a result of this change.

I start by considering the linear Fisher information (Eq. 5), and explicitly describe the summation over neurons:

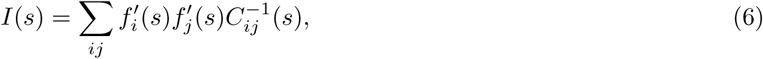

where the prime denotes a derivative with respect to the stimulus, and 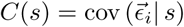 is the covariance matrix of the noise in the neural responses to stimulus *s*.

If neuron *k* were to be replaced by an untuned neuron, 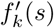 would be set to zero, and the population would now have a Fisher information value of

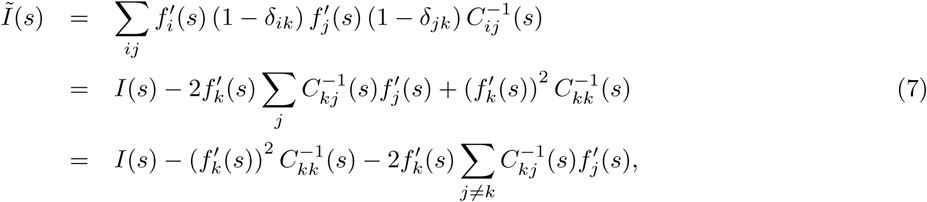

where *δ*_*ij*_ is the Kronecker delta (equal to 1 if *i* = *j*, and zero otherwise), and *I(s*) the Fisher information value from Eq. 6.

Whenever *Ĩ(s*) *> I(s*), the population code is made more informative by the inclusion of an untuned neuron. That condition corresponds to

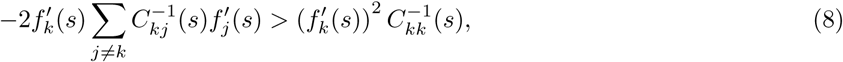

which is Eq. 1 of the main text.

## Analysis of *in vivo* neural recordings

### Overview of the experiment

The full description of the experiment is given by [13], and so I briefly summarize here. GCaMP6f was expressed in the excitatory neurons of the forebrain of mice. 2-photon imaging was used to measure the fluorescence of neurons in visual cortex through a cranial window. The mice were presented with drifting grating stimuli. The stimuli could move in any of 8 different directions, and at 6 different temporal frequencies. The stimuli were presented for 2 seconds each, followed by a 1 second gray screen before the next stimulus was presented. Each combination of direction and frequency was presented repeatedly (either 15 or 30 times each, depending on the temporal frequency).

## Data access

Following the example Jupyter notebook provided by [13] – which provides a template for accessing the experimental data – I retrieved the following data: average Δ*F/F* values for each neuron on each trial, and the stimulus direction for each trial. I analyzed all of the neurons observed in each experiment, and not only those that were labelled as visually responsive.

### Tuning curves

I calculated the tuning curves (Figs. 5A and B) by averaging the Δ*F/F* values for all trials of each direction: this *marginalizes* over the different temporal frequencies. The noise correlations coefficients (Fig. 5D) were computed over repeats of the same stimulus (same orientation and temporal frequency), and then averaged over all stimuli.

### Statistical tests of significance

For the results in Figs. 6, S4, and S5, I used two different methods to assess statistical significance. First, I used the standard paired sample t-tests. Second, I used non-parametric tests described below. These are typically more conservative because they are not sensitive to non-Gaussianity in the data. Both methods showed that the experimental results are significant.

For the non-parametric tests, I identified the number, *K*, of experiments in which the effect was positive. For example, for Fig. 6A, this was the number of experiments in which the full population gave better decoding performance than did the subset containing only tuned neurons. I then computed the probability that, of the *N* = 46 experiments, *K* or more of them would have a positive effect, if the outcome of each experiment came from an unbiased coin flip (which assigned a positive, or negative, effect with equal probability). This probability is obtained from the binomial distribution, and gives the probability that we would have observed the same results by random chance.

### Identifying tuned vs untuned neurons

Following [43, 9], direction selective cells were identified via their *circular variance*, with the direction selectivity index (DSI) computed for each neuron as follows. For each stimulus direction, I computed the two-dimensional direction vector *d(θ*) = [cos(*θ*), sin(*θ*)], and multiplied that by the neuron’s mean response to this stimulus *r(θ*) (i.e., the tuning curve value for that stimulus). This yielded a vector *v(θ*) = *r(θ)d(θ*) that points in the direction of the stimulus, with the length determined by the cell’s mean response to the stimulus. I then averaged this vector over all stimulus directions. If the neuron gave equal responses to all stimuli, the horizontal and vertical components of *v(θ*) would average out to zero over all the stimuli, whereas if the neuron responds selectively to one stimulus direction, this cancellation would not occur. Consequently, the DSI is measured by the length of ⟨*v(θ*)⟩, relative to the neuron’s mean response (averaged over all stimuli):

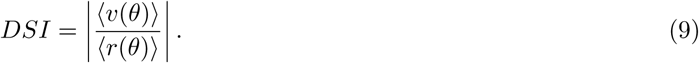

If the neuron responds strongly to only one stimulus direction, the DSI can be as large as 1, and the DSI can be as small as 0 for neurons that respond equally to all stimuli.

To identify tuned (vs “untuned”) neurons, I chose a cutoff of *DSI >* 0.25. This matches the smallest DSI of the direction-selective retinal ganglion cells studied by [9]. I also repeated the decoding analysis from Fig. 6 with different cutoffs on the DSI, and found qualitatively similar results: the precise value of the cutoff is unimportant (Figs. S4 and S5).

### Logistic regression decoding analysis

I used the logistic regression method of [44]. The classifier was trained to take in vectors of neural responses, in response to two different stimuli, and to return labels (“0” or “1”) that indicate which of the two stimuli was presented on each trial. I randomly divided the data into a training set (75% of the data) that was used to fit the weights of the classifier, and a test set (25% of the data) that was used to measure the performance. After training on the training data, I applied the classifier to the neural responses from the test data set, yielding an output value for each response vector. Values above 0.5 indicated that the stimulus was most likely to be stimulus “1”, whereas values less than 0.5 were taken to indicate that response was most likely generated by stimulus “0”. I then computed the fraction of these test trials on which this classifier correctly identified the stimulus that caused the neural response. This analysis was separately done for all (8 ×; 7)/2 = 28 different stimulus pairings: reported performance values are averages over all such stimulus pairings.

I trained the classifier using Ridge regression. This method penalizes the classifier for having overly-strong weights, and helps improve generalization performance. I chose the value of the regularization penalty, *λ*_*reg*_, to be around the saturation point of the performance vs. *λ*_*reg*_ curve (Fig. S6).

## Acknowledgments

Thanks to Eric Shea-Brown and Alex Cayco-Gajic for helpful discussions, and to the Allen Institute for sharing their *in vivo* datasets. JZ gratefully acknowledges the following funding: Canadian Institute for Advanced Research (CIFAR) Azrieli Global Scholar Award, Sloan Research Fellowship, and Google Faculty Research Award.

## Supplemental Information and Figures

**Figure S1: different parameters for the limited-range correlations**

**Supplementary Figure 1:**
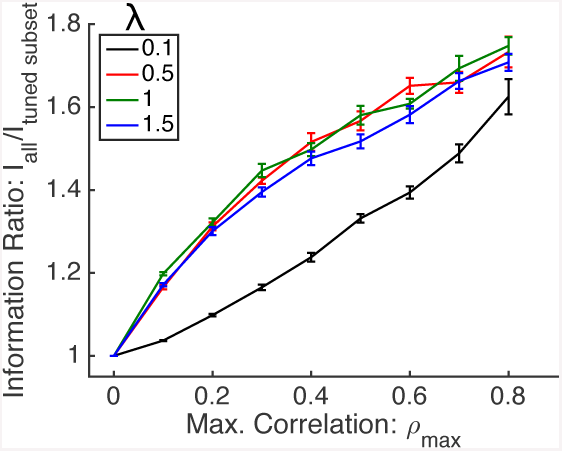
Dependence of information on limited range correlation parameters. (Related to Fig. 1.) I repeated the calculations from Fig. 1, in all cases for populations of 200 neurons. I repeated the calculations for different values of *ρ*_*max*_ and *λ*, the parameters that define the limited-range correlations. For each set of parameters, I computed the ratio of Fisher information in the full population of 200 neurons, vs. the Fisher information in just the tuned subset of (70% of) the population. Error bars are the S.E.M. over 10 random sets of different tuning curves.

### Information-limiting correlations and Figs. S2 and S3

I repeated the calculations from Fig. 3, using the same random tuning curve shapes (as in Fig. 1A), but different covariance matrices. Specifically, in Fig. S2, the model neurons had the *differential correlations* studied by [19, 23]. These correlations are such that the shared (correlated) part of the population noise mimics the changes in neural firing pattern induced by changes in the stimulus, thereby causing the distributions of responses to different stimuli to overlap substantially. As a a result of that overlap in the stimulus-evoked response distributions, the noise substantially hinders the population code, and thus information saturates with increasing population size.

That covariance matrix, *C*_*diff*_, is given by

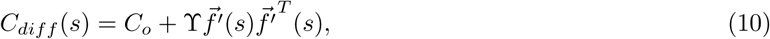

where ϒ is a (small) scalar parameter that sets the strength of the differential correlations, and *C*_*o*_ is a covariance matrix that does not lead to information saturating with population size. For this analysis, I chose *C*_*o*_ to be the covariance matrix corresponding to neurons with limited range correlations and Poisson variability (e.g., for ? = 0, this is the same covariance matrix as in Fig. 3). For the results in Fig. S2, I chose ϒ = 1×10^−3^, corresponding to weak but non-zero differential correlations; I later consider the case of stronger differential correlations. (Note that, with information-limiting correlations, the Fisher information saturates at ϒ^−1^. Moreover, cell pairs containing untuned neurons always have zero entries in the 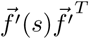 matrix; any correlations for cell pairs containing untuned neurons come from the *C*_*o*_ component of the covariance matrix.)

For Fig. S2A, the covariance matrix *C*_*o*_ matches the one from Fig. 3A; cell pairs containing untuned neurons have no noise correlations. For Fig S2B, covariance matrix *C*_*o*_ matches the one from Fig. 3B; cell pairs containing untuned neurons do have noise correlations.

For the tuned subset of the population, the noise structure in Fig. S2B was identical to the one in Fig. S2A (both contain differential correlations and limited-range correlations), and thus the tuned subsets of neurons have the same information in both cases. (Red data points in Fig. S2A have the same values as do points on the red curve in Fig. S2B).

**Supplementary Figure 2:**
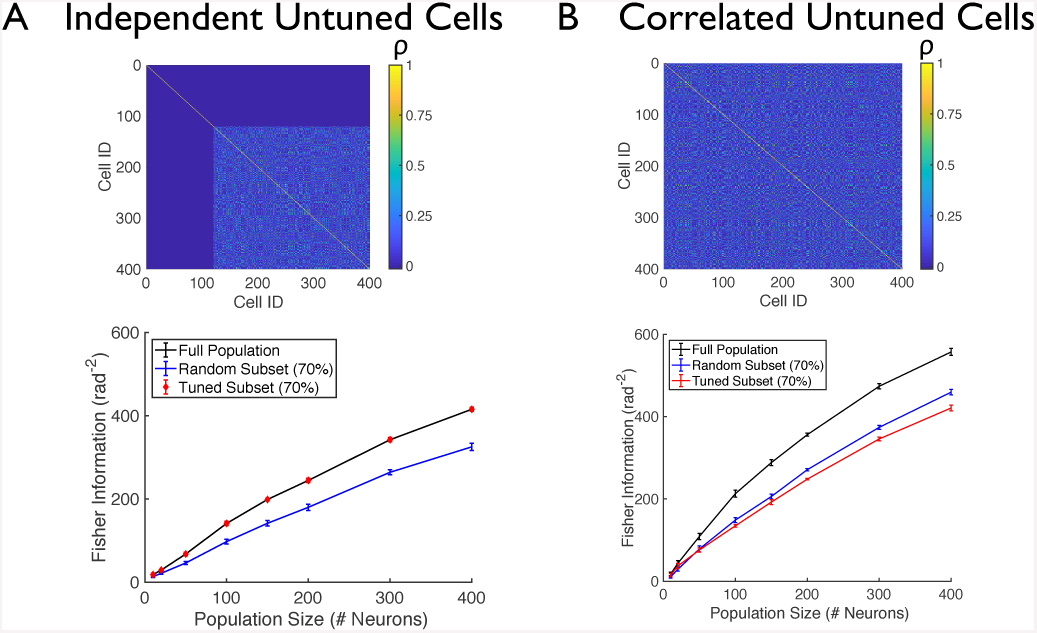
Untuned neurons improve population coding when they are correlated with the tuned neurons. (Similar to Fig. 3, but with information-limiting correlations) I considered neural populations with tuning curves as in Fig. 1, and where the untuned neurons were either independent of the tuned ones (A), or where the untuned neurons were correlated with the tuned ones (B). 70% of the neurons in each population were tuned to the stimulus, and 30% were untuned. Upper panels show correlation matrices for 400-cell populations: cells 1 through 120 are untuned, whereas the remainder were tuned. Bottom panels show the Fisher information for the full neural populations (black), for the tuned subsets of neurons (red), and for random subsets of 70% of the neurons in each population (blue). (Data points shown are mean ± S.E.M., computed over 5 different random draws of the tuning curves)

Different from Fig. S2A, the full population in Fig. S2B contained much more information than did the tuned subset, indicating that the untuned neurons do contribute to the population code in this case. Thus, the observation that, when there are correlations between tuned an untuned neurons, untuned neurons improve population coding, holds in the case of information-limiting correlations (Fig. S2).

To determine how the effects seen in Fig. S2 depend on the strength of the differential correlations, I repeated those calculations, using a value of ϒ = 5 ×; 10^−3^. In the case, the information saturates at 200 rad^−2^. The results of those calculations show that, even in the case of stronger differential correlations, the untuned neurons can improve the population code, so long as they are correlated with the other neurons (Fig. S3). Intuitively, the untuned neurons help the population reach the point of saturating information more quickly, even if they cannot enable the population to surpass the ϒ^−1^ limit on information set by the differential correlations.

### Varying the DSI cutoff for labelling cells as “tuned” vs “untuned”; Figs. S4 and S5

I repeated the analysis from Fig. 6 with two different cutoffs on the direction selectivity index (DSI), which is used to distinguish “tuned” neurons from “untuned” ones.

<u>DSI cutoff of 0.3 (Fig. S4):</u>

I labeled cells with *DSI >* 0.3 as “tuned” and those with *DSI <* 0.3 as “untuned”. I then compared the logistic regression decoding performance on the full population with that on the tuned subset of the population. The full population yielded 13 ± 2% better decoding performance (*p* = 4.7 ×; 10^−12^, paired sample single-sided t test; *p <* 10^−8^, non-parametric binomial test of significance).

**Supplementary Figure 3:**
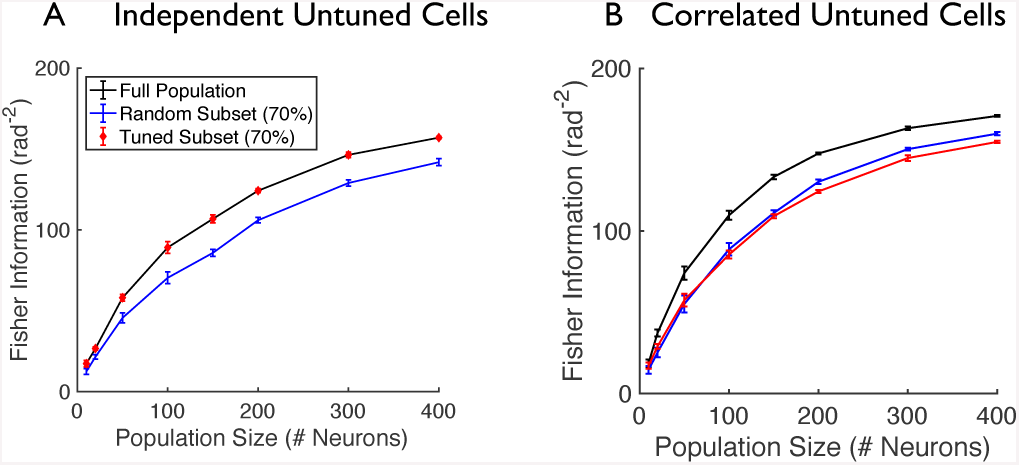
Untuned neurons improve population coding when they are correlated with the tuned neurons. (Same as Fig. S2 lower panels, but with stronger differential correlations) I considered neural populations with tuning curves as in Fig. 1, and where the untuned neurons were either independent of the tuned ones (A), or where the untuned neurons were correlated with the tuned ones (B). 70% of the neurons in each population were tuned to the stimulus, and 30% were untuned. Bottom panels show the Fisher information for the full neural populations (black), for the tuned subsets of neurons (red), and for random subsets of 70% of the neurons in each population (blue). (Data points shown are mean ± S.E.M., computed over 5 different random draws of the tuning curves).

**Supplementary Figure 4:**
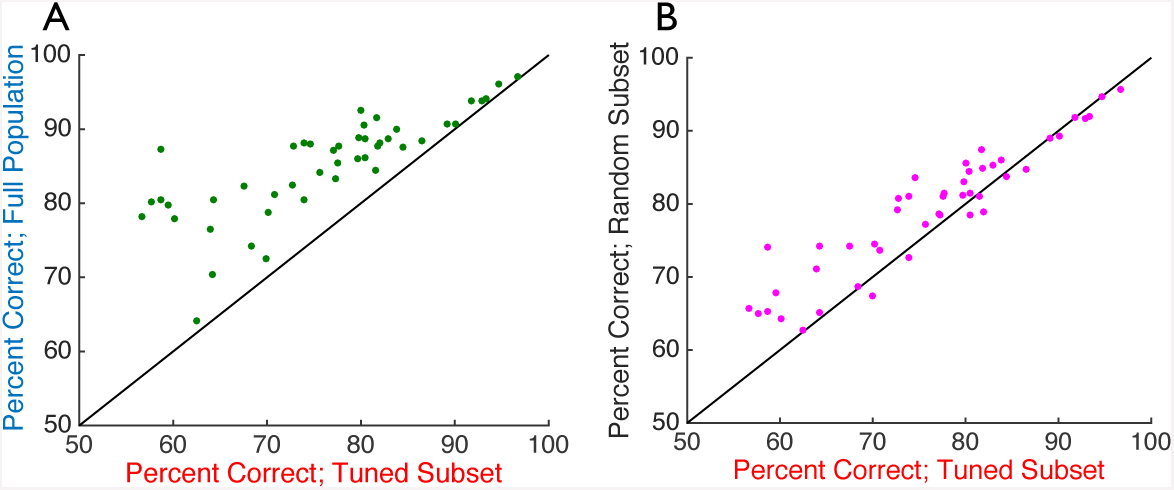
Untuned neurons can enhance information coding in *in vivo* neural populations. (Similar to Fig. 6, but with a cutoff of *DSI >* 0.3 for labeling cells as “tuned”.) I used logistic regression to perform pairwise discrimination on the population response vectors, to determine which of 2 different stimuli caused each response. I repeated this analysis for all possible pairs of stimuli: reported values are the percentage of trials for which the stimulus was correctly identified, averaged over all possible pairings (there is one data point per experiment). (A) Decoding accuracy when the full population response vectors were decoded (vertical axis) vs. when only the tuned subsets of the neurons are seen by the decoder (horizontal axis). (B) Decoding accuracy when random subsets of the neurons (of the same size as the tuned subset, but containing both tuned and untuned neurons) are input to the decoder (vertical axis) vs. when only the tuned subsets of the neurons are seen by the decoder (horizontal axis). Chance performance for this binary discrimination task is 50%. Diagonal line denotes equality.

Next, I asked whether populations that include both tuned and untuned neurons could yield better decoding vs populations of the same size but containing only tuned cells. To answer this question, I extracted a random subset of the neurons from each population, that was the same size as the set of tuned neurons within that population. I then performed the logistic regression analysis on these random subsets, and compared the performance with that which was obtained on the tuned subsets. On average, the decoding performance was 5 ± 1% (mean ± S.E.M.) better using the random subsets vs the fully tuned ones, a modest but statistically significant difference (*p* = 3.9 ×; 10^−6^, paired single-sided t-test; and *p* = 2.2 ×; 10^−3^, non-parametric binomial test of significance).

<u>DSI cutoff of 0.2 (Fig. S5):</u>

I labeled cells with the *DSI >* 0.2 as “tuned” and those with *DSI <* 0.2 as “untuned”. I then compared logistic regression decoding performance on the full population with that on the tuned subset of the population. The full population yielded 7 ± 1% better decoding performance (*p* = 9.8 ×; 10^−10^, paired sample single-sided t test; *p <* 10^−8^, non-parametric binomial test of significance).

**Supplementary Figure 5:**
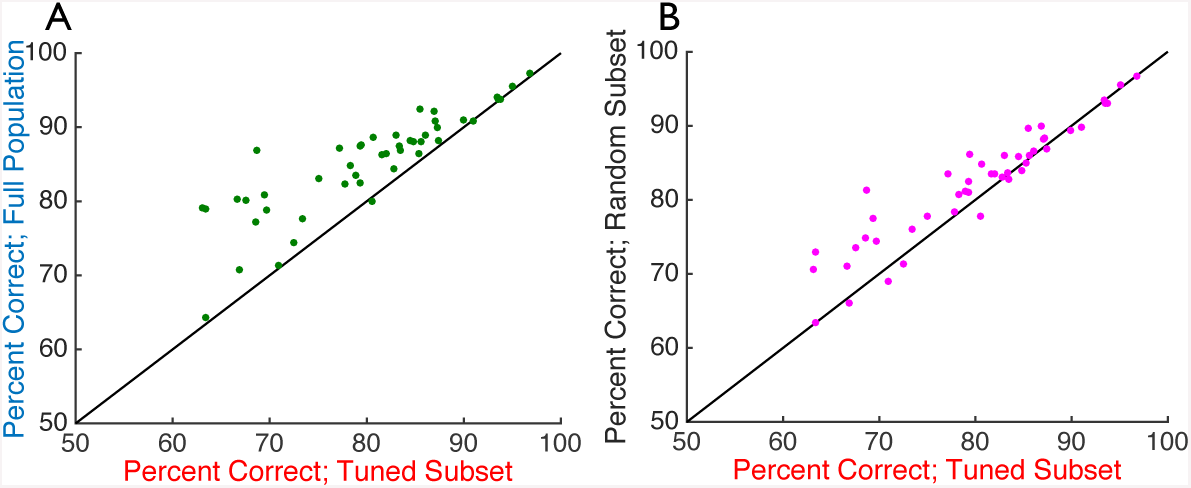
Untuned neurons can enhance information coding in *in vivo* neural populations. (Similar to Fig. 6, but with a cutoff of *DSI >* 0.2 for labeling cells as “tuned”.) I used logistic regression to perform pairwise discrimination on the population response vectors, to determine which of 2 different stimuli caused each response. I repeated this analysis for all possible pairs of stimuli: reported values are the percentage of trials for which the stimulus was correctly identified, averaged over all possible pairings (there is one data point per experiment). (A) Decoding accuracy when the full population response vectors were decoded (vertical axis) vs. when only the tuned subsets of the neurons are seen by the decoder (horizontal axis). (B) Decoding accuracy when random subsets of the neurons (of the same size as the tuned subset, but containing both tuned and untuned neurons) are input to the decoder (vertical axis) vs. when only the tuned subsets of the neurons are seen by the decoder (horizontal axis). Chance performance for this binary discrimination task is 50%. Diagonal line denotes equality.

Next, I asked whether populations that include both tuned and untuned neurons could yield better decoding vs populations of the same size but containing only tuned cells. To answer this question, I extracted a random subset of the neurons from each population, that was the same size as the set of tuned neurons within that population. I then performed the logistic regression analysis on these random subsets, and compared the performance with that which was obtained on the tuned subsets. On average, the decoding performance was 3 ± 1% (mean ± S.E.M.) better using the random subsets vs the fully tuned ones, a modest but statistically significant difference (*p* = 2.7 ×; 10^−5^, paired single-sided t-test; and *p* = 5.7 ×; 10^−3^, non-parametric binomial test of significance).

### Setting the value of the regularization parameter

I repeated the logistic regression analysis for many different values of the regularization parameter, *λ*_*reg*_, and assessed the classification performance (on the held-out test data) for each value. I then chose the value of *λ*_*reg*_ to be near the saturation point of the performance vs *λ*_*reg*_ curves (Fig. S6: vertical line). Data shown are performance when decoding the full populations. Similar curves were obtained when decoding random subsets, or only the tuned subsets of the population (not shown).

**Supplementary Figure 6:**
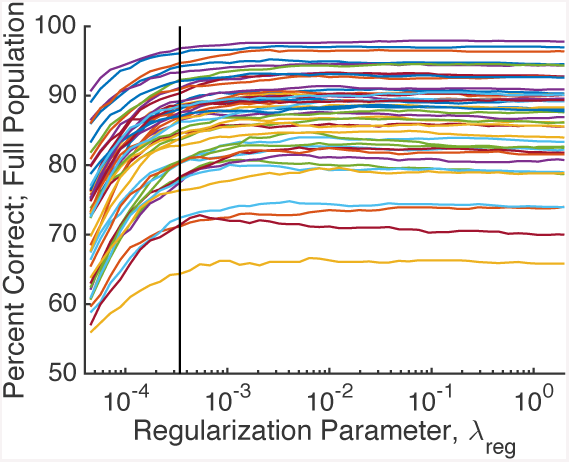
Setting the regularization parameter. Classification performance vs Ridge regularization parameter *λ*_*reg*_ used in training the logistic regression classifier, shown for each different experiment (each curve is for a different imaging experiment). Vertical line indicates the chosen *λ*_*reg*_ value, which was selected to fall near the saturation points of the curves.

